# The genomic impact of European colonization of the Americas

**DOI:** 10.1101/676437

**Authors:** Linda Ongaro, Marilia O. Scliar, Rodrigo Flores, Alessandro Raveane, Davide Marnetto, Stefania Sarno, Guido A. Gnecchi-Ruscone, Marta Alarcon-Riquelme, Etienne Patin, Pongsakorn Wangkumhang, Garrett Hellenthal, Miguel Gonzalez-Santos, Roy J. King, Anastasia Kouvatsi, Oleg Balanovsky, Elena Balanovska, Lubov Atramentova, Shahlo Turdikulova, Sarabjit Mastana, Damir Marjanovic, Lejla Kovacevic, Andreja Leskovac, Maria F. Lima-Costa, Alexandre C. Pereira, Mauricio L. Barreto, Bernardo L. Horta, Nédio Mabunda, Celia A. May, Andres Moreno-Estrada, Alessandro Achilli, Anna Olivieri, Ornella Semino, Kristiina Tambets, Toomas Kivisild, Donata Luiselli, Antonio Torroni, Cristian Capelli, Eduardo Tarazona-Santos, Mait Metspalu, Luca Pagani, Francesco Montinaro

## Abstract

The human genetic diversity of the Americas has been shaped by several events of gene flow that have continued since the Colonial Era and the Atlantic slave trade. Moreover, multiple waves of migration followed by local admixture occurred in the last two centuries, the impact of which has been largely unexplored.

Here we compiled a genome-wide dataset of ∼12,000 individuals from twelve American countries and ∼6,000 individuals from worldwide populations and applied haplotype-based methods to investigate how historical movements from outside the New World affected i) the genetic structure, ii) the admixture profile, iii) the demographic history and iv) sex-biased gene-flow dynamics, of the Americas.

We revealed a high degree of complexity underlying the genetic contribution of European and African populations in North and South America, from both geographic and temporal perspectives, identifying previously unreported sources related to Italy, the Middle East and to specific regions of Africa.

## Introduction

North and South America were the last two continents to be colonized by humans. The peopling of the Americas was a complex process, involving multiple dispersal events, that started at least 15 thousand years ago (kya) ^1–6^. Nowadays, a substantial proportion of individuals living in the Americas is the result of more recent episodes of admixture, occurred following large migrations during and after the European Colonial era and the consequent deportations in the African slave trade ^7^.

The Colonial Era of the Americas started soon after the European discovery of the continents in 1492, when old world’s powers started to explore and settle the Western hemisphere. This colonization heavily impacted autochthonous population, which were decimated both by wars and pathogens brought by the invaders. The Atlantic slave trade, which occurred between the 16^th^ and 19^th^ century, was initiated by Portuguese and Spaniards leading to the presence of millions of people with African ancestry in the American continents.

Historical records have attested a general imbalance in the number of males and females disembarked in these migration and deportation events. Especially during the early phase of Iberian colonization, the immigrants were represented mostly (>80%) by males ^8^, while the females represented only 5-6%, although their proportion increased in the following decades^7^. Since the end of the 19^th^ century, several migrations, mostly from the Southern and Eastern regions of Europe, had a strong impact on the demographic variability of the continent. In fact, it has been estimated that more than 32 million individuals reached the United States at the end of the 1800s and the beginning of the 1900s and similar estimates are available for other American countries. For example, more than 6 million people arrived in Argentina and more than 5 million in Brazil in the same period ^9^.

Given their historical and epidemiological implications, these migrations have been the subject of several genetic studies ^10–14^. Most of them have exploited Local Ancestry inference (LA) algorithms, in which individual genomes are deconvoluted into fragments ultimately tracing their ancestry to populations from different macro-geographic areas. LA approaches provided multiple insights into the composition of several recently admixed populations ^15,16^. However, when multiple closely related populations are involved in the admixture of a specific target group, this strategy might have a reduced power in discriminating among sources, leading to spurious or incomplete results.

While several surveys ^10,13,14,17^ present a continental-wide analysis of the origin and dynamics of the African and European Diaspora into the Americas, a more comprehensive and systematic investigation considering multiple ancestries across the two continents is currently missing ^10–13^.

Gouveia et al.^17^ have recently performed a detailed analysis of the African regional ancestry and its dynamics in several populations from North-, South-America and the Caribbean region.

The recently increased availability of genome-wide data, offers, for the first time, the chance to capture the complexity of historical and demographic events that affected the recent history of the Americas by studying the recent admixture profile of American populations in the continents.

With this in mind, we have assembled and analysed a genome-wide dataset of 17,722 individuals, including ∼12,000 from North, Central and South America and ∼6,000 from Africa, Europe, Asia and Oceania (Supplementary Figure 1, Supplementary Table 1A-B).

To provide a comprehensive genetic description of the complex ancestries blending in the Americas, we have harnessed haplotype-based and allele frequency methods to a) reconstruct the fine scale ancestry composition, b) evaluate the time of admixture, c) explore the demographic evolution of different continental ancestries after the admixture and d) assess the extent and magnitude of sex biased gene-flow dynamics.

## Results

### Clustering of the donor individuals

To minimize the impact of within-source (“donors”) genetic heterogeneity in the ancestry characterization process, we grouped the assembled 6,115 individuals (Supplementary Figure 1, Supplementary Table 1A-B) from 239 population-label donors (from which American individuals are subsequently allowed to copy fragments of genome, see Methods) into 89 genetically homogeneous clusters (Supplementary Figure 2, Supplementary Table 2) on the basis of haplotype similarities using CHROMOPAINTER and fineSTRUCTURE^18^. 41 and 40 of these were European/West Eurasian and African respectively, along with 3 groups of American individuals; while the remaining 5 clusters differentiate Oceania and East Asia. A detailed description of the composition of the clusters is reported in Supplementary Text and Supplementary Table 2.

Our fineSTRUCTURE results (Supplementary Figure 2, Supplementary Table 2) confirm the worldwide genetic variation pattern already observed by previous studies at the continental scale ^19–23^.

### The ancestral mosaic of American populations

We fit each of the 22 American populations as a mixture of the identified donor groups using SOURCEFIND^24^. In contrast to Non-Negative Least Square (NNLS) approach, SOURCEFIND uses a Bayesian algorithm to provide increased resolution in distinguishing true contribution from background noise (see Methods section).

The contribution of the 21 most representative clusters (sources with proportion of no less than 2% in at least one recipient population) to the American admixed populations are reported in Figure 1A and Supplementary Table 3. The same procedure using NNLS provided consistently similar results (Supplementary Figure 3).

**Figure 1.**
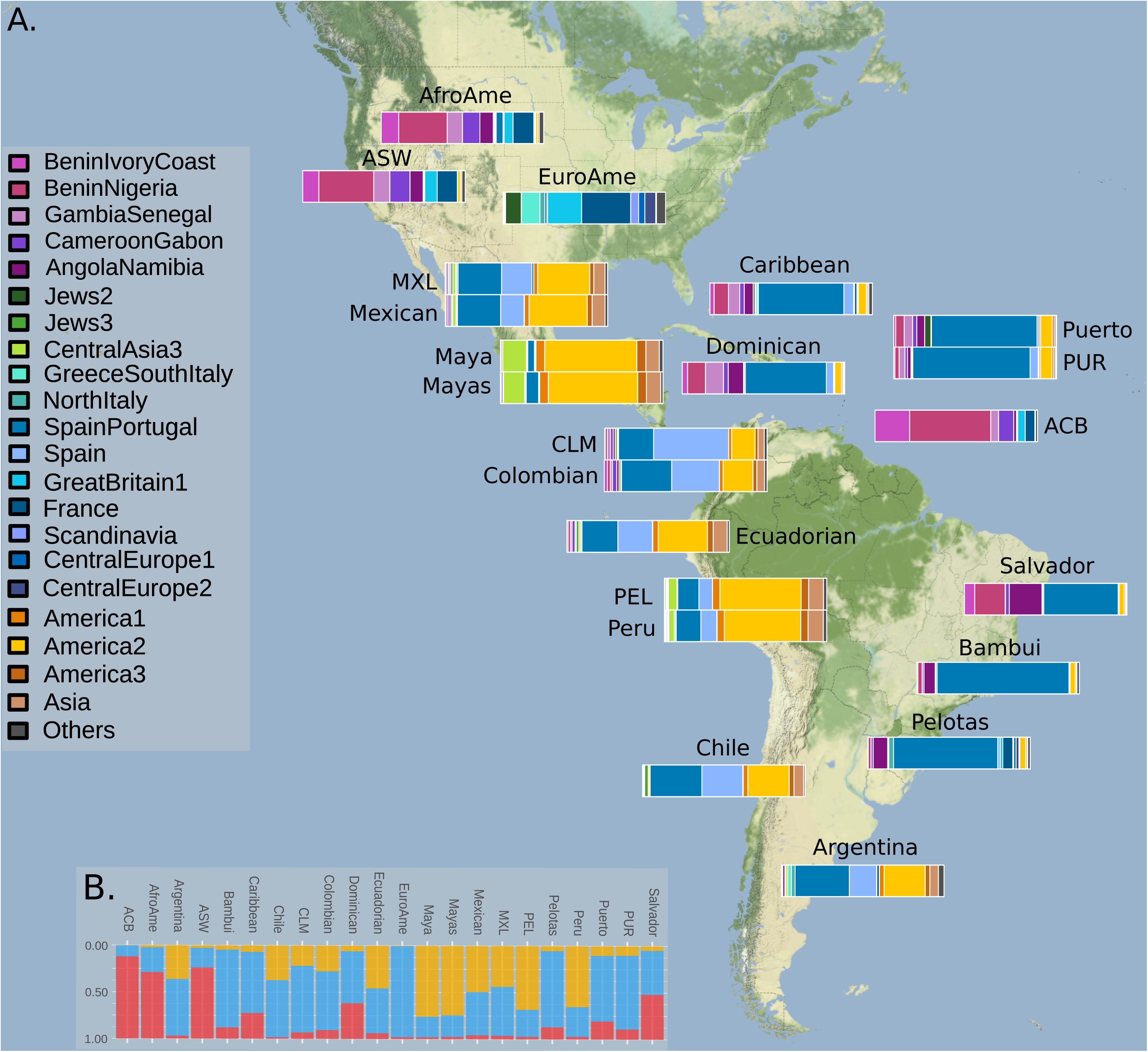
The ancestral mosaic of American populations reveals a highly complex ancestral composition. **A)** Barplots representing ancestral genetic proportions based on SOURCEFIND results for North and South American populations. We applied CHROMOPAINTER/fineSTRUCTURE and SOURCEFIND to find the ancestral compositions of 22 American populations. Only the contribution for the 21 most representative fineSTRUCTURE clusters (contributing ≥ 2% in at least one recipient population) is reported (Supplementary Table 3). **B)** Proportion of continental ancestries for all target populations. Ancestries are represented in red for Africa, blue for Europe and yellow for America/Asia.

#### African ancestries distribution reflects the complexity of the Slave Trade dynamics

Sub-Saharan African ancestry was observed at high proportion in African-Americans (AfroAme: 69% and ASW: 74.1%) and Barbados (ACB: 87.1%), with relatively high contribution registered also for the other Caribbean and Brazilian populations (>10%; Figure 1B).

In detail, “*BeninNigeria*” cluster showed the highest contribution (≥30% of the total) in African Americans and Barbados, while, in other Caribbean populations, the contribution of “*BeninNigeria*” and “*GambiaSenegal*” clusters is comparable with average proportion of 6.9% (min=2.6%; max=11%) and 6.7% (min=3.6%; max=11.1%), respectively.

Moreover, we found contributions from “*GambiaSenegal*” (mean=4.2%; min=1.3%; max=11.1%) in Mexico, Caribbean islands and Colombia but not in Brazil, Argentina and Chile that have a proportion of less than 0.2%, consistent with previous results^17^.

In South America, all the analysed populations show high heterogeneity in African proportions, the highest values in individuals from Salvador (47.8%)^25^, possibly reflecting the high number of deported African slaves for sugar production in the Northeast area of Brazil in the 17^th^ century ^26^.

In details, the African cluster contributing the most is related to groups from Angola and Namibia (*“AngolaNamibia”* cluster), with Salvador (Brazil) having the highest percentage (>20%), similar to the contribution from “*BeninNigeria”* (∼19%), mirroring the history of African slaves arrivals in Brazil ^26^ (Supplementary Figure 4, Supplementary Table 3). Although a non-negligible contribution from East and South-East Africa at the end of the Slave Trade period has been documented ^27^, none of the analysed population samples showed an East African ancestry fraction larger than 2%. AfroAme and ASW show the highest proportion of this ancestry (1.2% and 0.8%, respectively). Nevertheless, when the ancestry is explored at individual level, samples with more than 5% of East and/or South-East African ancestries were present in more than 1% of individuals from AfroAme (30/2004), ASW (2/55), Bambui (10/909) and Pelotas (51/3629) (Supplementary Figure 5), supporting recent findings ^17^.

When dissecting the African ancestry into regional sources (Supplementary Figure 6B), the UPGMA clustering does not strictly mirror geographical/historical patterns. Yet, all the Caribbean and circum-Caribbean populations, with the exception of a Colombian sample, cluster together. Similarly, all the Southern American samples, but not Chile, form a private group. Interestingly, ACB is different from any other populations, composed mainly by “*BeninIvoryCoast*” and “*BeninNigeria*” clusters.

#### Complex variation of European ancestries distribution

European ancestry was observed at high proportion in European-Americans (EuroAme), Caribbean Islands (PUR from Puerto Rico having the highest proportion, 79%) and Mexico (∼42% and ∼48% for Mexican and MXL, respectively), but also in Southern America, with proportions ranging from 22% in Peru (PEL) to ∼82% in Bambui.

When the variation of European ancestries in the Americas is evaluated groups from United States (EuroAme, AfroAme e ASW) and Barbados (ACB) are characterized by a substantial proportion of British and French ancestries. On the contrary, in the remaining populations the most prominent European ancestry was represented by Iberian-related clusters, reflecting the geo-political extent of European occupation during the Colonial Era (Figure 1A). In details, populations from Mexico, Caribbeans and South America derive most of their European ancestry from the Iberian Peninsula, represented by two clusters. European Americans (EuroAme) exhibit high levels of heterogeneity, showing not only a high proportion of France and Great Britain, but also Greece and South Italy, Central Europe and Scandinavia, revealing the high variability of European ancestries in the United States, possibly due to secondary movements in the 19^th^ and 20^th^ centuries ^28^, which involved populations that did not take part in the Colonial Era movements^9^. Moreover, Pelotas (Brazil) is characterized by a high contribution from North Italy (∼3%), while Argentina from both North and South Italy (2.3% and 2.2%, respectively).

The investigation of the individual ancestry profiles confirmed and further refined the identification of multiple European secondary sources.

In one African American sample (AfroAme), we identified a high variability of European ancestry, with several individuals characterized by more than 5% ancestry from Northern, Central and Southern European regions (Supplementary Figure 5).

Italian ancestry was also found at considerable proportion (>5%) in individuals from Colombia (4/98), Caribbean (51/1112), Dominican Republic (2/27), Ecuador (1/19), Mexico (15/427), Peru (6/153), Puerto Rico (4/99), Argentina (27/133) and Brazil (622/5779). In fact, Italy has been reported as one of the main sources of migrants to South America during the 19th century, second only to the Spanish and Portuguese influences ^29^ (Supplementary Figure 5, Supplementary Figure 7).

We estimated the relationship among American populations considering the relative European ancestries proportion by applying a UPGMA clustering approach (Supplementary Figure 6A). Differences in regional affinities to British/French vs Spanish/Portuguese ancestries among American populations were observed. Furthermore, within the last group, Spanish and Portuguese ancestries show distinct geographical distributions, consistent with the Treaty of Tordesillas, signed in 1494, to regulate the regional influence of Spain and Portugal in the Americas (Caribbean islands represent again an exception) (Supplementary Figure 6A).

#### Native American ancestry distribution

With the exception of Mayan individuals (>65%), Native American ancestry is high in populations from the Southern part of the continent and Mexico (41%), with the highest values in Peru (59.2% PEL), Ecuador (37%) and Argentina (31%) (Figure 1 and Supplementary Table 3). Interestingly, in both the analysed African-American samples we identified a non-negligible proportion of individuals harbouring Native American related ancestry.

#### The contribution of Jewish related ancestry in the Americas

A recent genetic investigation found a non-negligible proportion of ancestry related to Jews and Middle East groups in five populations from Southern America (Mexico, Colombia, Peru, Chile, Brazil) ^24^. In our analysis, we confirmed the presence of genetic ancestries related to “*NorthAfrica*”, “*Levant*”, “*LevantCaucasus*” and “*Jews*” clusters in the same countries, although at a lower proportion than previously estimated (∼2.8%). This discrepancy might be due, at least in part, to the fact that our dataset is mostly composed by Brazilian individuals, which have been documented to have a smaller Jewish ancestry ^24^. Only 2.5% of analysed individuals contain more than 5% of Jewish or Middle-Eastern ancestry (Salvador: 0.8%, Bambui: 3.2%, Pelotas: 2.9 %). In contrast, this proportion is higher in the non-Brazilian populations (Colombian from Medellin CLM: 8%, Colombian: 3.8%, Peru: 2.3%, Mexican: 5.4%, Mexicans from Lima, MXL: 11%, Chile: 16%, Argentina 12%). Similar proportions were found for Caribbean populations (ACB from Barbados: 1.4%, Caribbean: 6.8%, Dominican: 3.7%, Puerto: 3.9%, PUR: 1.4%). Interestingly, we found a relatively high proportion of individuals showing more than 5% contributions related to “Jewish” sources also in one of the African American samples (AfroAme: 3.8%), in European Americans (EuroAme: 26.7%) and in Argentinians (∼12%) (Supplementary Figure 5).

### Assessing the impact of sex-biased admixture of the Americas

To evaluate the impact of sex-biased admixture dynamics in the American populations, we compared the continental ancestry proportions inferred by ADMIXTURE ^30^ from autosomal data against those estimated for the X chromosome (see Methods). With respect to European ancestry, a paired Wilcoxon test comparing the distribution of autosomal vs X chromosome revealed that the former is significantly higher in all comparisons, suggesting a higher contribution of European males than females in the gene pool of American populations (Supplementary Figure 8, Supplementary Table 4), in agreement with previous continental-scale reports based on more limited data ^25,31,32^. This observation is further supported by the fact that Native American ancestry estimated from autosomal data is always lower (with the exception of Dominican) than that estimated from the X chromosome. In contrast, when considering the African ancestry, a considerable number of populations do not show any signature of sex imbalance. Indeed, in only eight out of 19 comparisons (ACB, AfroAme, Bambui, Caribbean, EuroAme, Pelotas, PUR and Salvador) the autosomal proportion was significantly lower than that inferred from the X chromosome (adj. p < 0.05). With the exception of ACB, all these significant differences were associated with sample sizes greater than 100. These results are in contrast with historical records documenting a higher number of disembarked male slaves ^27^ and might reflect complex admixture dynamics in the following five centuries, or limitations in the approach exploited here, as previously suggested ^33^.

### Inferring the time of admixture in American populations

To provide a temporal dimension to the gene flow among the analysed populations, we inferred time of admixture by applying GLOBETROTTER (GT) in two different setups for “*Population”* and *“Individual”* level analyses, as detailed in the Methods section.

In population-level inferences all the analysed groups showed evidence of at least one admixture event as reported in Figure 2A and Supplementary Table 5. Specifically, we identified one admixture event in 14 populations (ASW, ACB, Mayas, Maya, PEL, Peru, Salvador, Ecuadorian, Colombian, MXL, Argentina, CLM, Chile, Puerto) with inferred times spanning between ∼6 and 11 generations ago. The identified sources are related to British or French and Benin-Nigeria in ACB and ASW, Iberian or Southern European and America in Maya, Mayas, PEL, Puerto, Peru, Ecuador, Colombian, CLM, MXL, Argentina and Chile, in line with SOURCEFIND estimates. In contrast, Salvador sources are representative of Iberia and Cameroon-Gabon. Two populations from Caribbean Islands, PUR and Dominican, showed a curve profile that fits better with a single admixture involving more than two sources from Europe, Africa and America, dated ∼9-11 generations ago. The remaining six populations (Mexican, EuroAme, Pelotas, Caribbean, AfroAme and Bambui) showed signature of at least two admixture events mainly involving American, European and African sources and occurring 6-8 generations ago.

**Figure 2.**
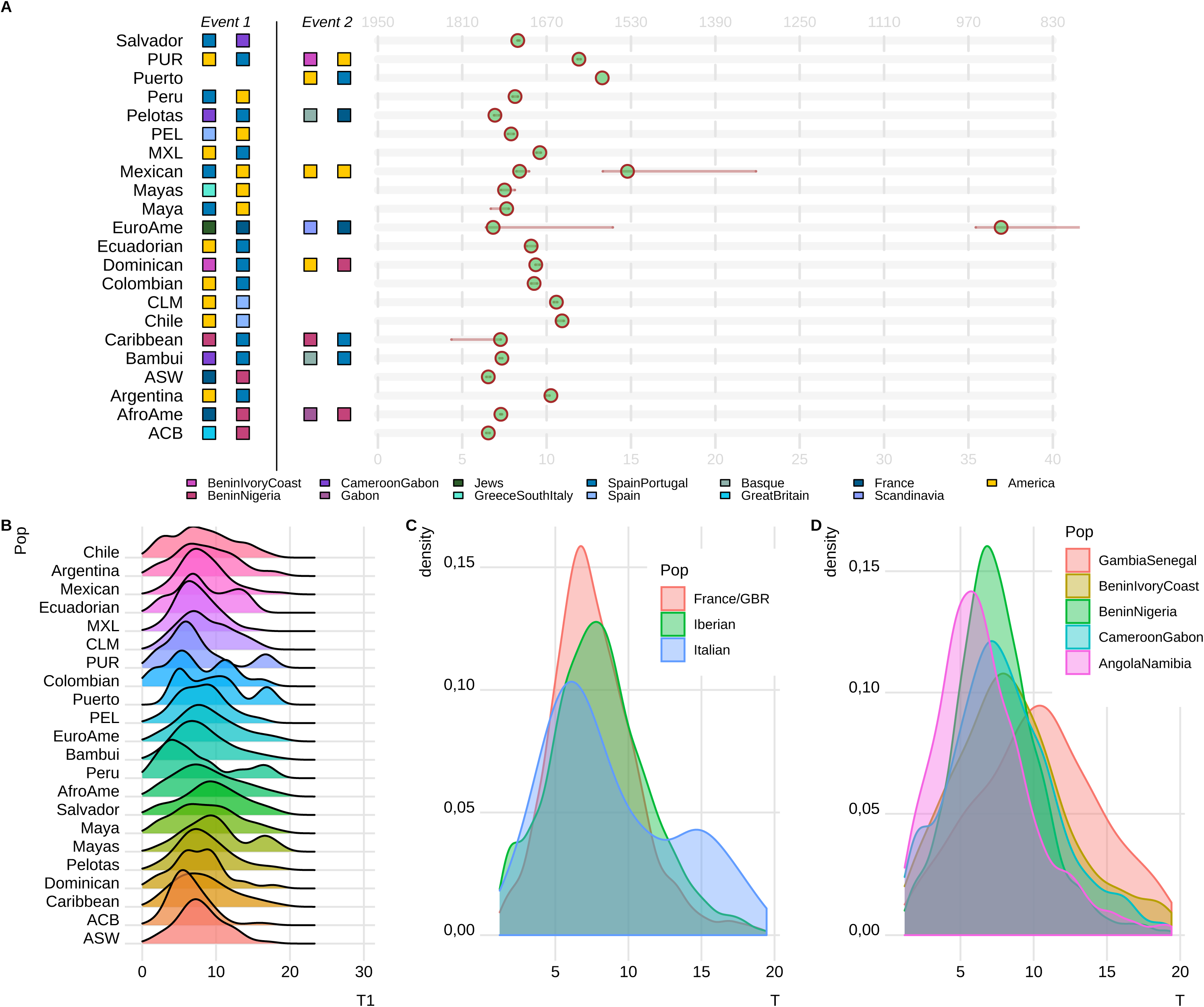
The admixture history of the Americas, as inferred by GLOBETROTTER (GT). **A)** Estimates of time and sources of admixture events considering the whole population as target. One or two events of admixture are reported for each population. The closest inferred sources of admixture, are represented as colored squares, circles show the corresponding time of admixture estimated by GT. Time is expressed in generations from present (bottom x axis), and years of CE (top x axis). **B)** Distribution of admixture times considering single individuals as targets. We retained only the 2.5%-97.5% distribution of time estimation for each population. **C)** Density of admixture times inferred in events considering France/GBR, Iberian, and Italian clusters as sources, for all the 11,607 admixed American individuals under study. **D)** Density of admixture times inferred in events considering “*GambiaSenegal”*, “*BeninIvoryCoast”*, “*BeninNigeria”*, “*CameroonGabon*”, “*Gabon”* and “*AngolaNamibia”* clusters as sources, for all the 11,607 admixed American individuals under study.

To assess regional spatio-temporal differences in admixture dynamics, we performed a GT *“Individual”* analysis (Figure 2B-D). For all the analysed populations the inferred 2.5%-97.5% time interval had similar boundaries spanning between ∼1 and ∼20 generations ago (min=1.18; max=19.5).

The source-specific admixture time estimates were explored evaluating the distributions of time inferred considering different European and African signals (Figure 2B-C). When the European sources were considered, times involving Iberian clusters were significantly older than those involving British/French ones, which in turn were characterized by dates significantly older than those involving Italian sources (Wilcoxon test, Bonferroni adjusted p-value < 0.05).

For the five African sources considered, times inferred for the “*SenegalGambia”* cluster are significantly older than all the other tested sources (Wilcoxon test, Bonferroni adjusted p-value < 0.05). In contrast, times involving “*AngolaNamibia”* are more recent than all the others (Wilcoxon test, Bonferroni adjusted p-value < 0.05). Moreover, times involving “*BeninIvoryCoast”* are significantly older than the one involving “*BeninNigeria*” and “*CameroonGabon*”. Lastly, times involving “*CameroonGabon”* are significantly older than the one involving “*BeninNigeria”* (Wilcoxon test, p-value < 0.05).

### Reconstructing the ancestry specific demographic histories of admixed populations

To characterize the demographic history of specific continental ancestries, we intersected the results of Identity-By-Descent (IBD) and LA inferences as in Browning et al. ^34^. We excluded from the analysis all the population ancestries in which where *α* (*continent*) * *N* < 50 where α is the proportion of a specific ancestry as estimated by SOURCEFIND, and N is the total number of chromosomes in the analysed population.

The majority of the studied populations showed, for all the continental ancestries considered, a demographic curve characterized by a decline until approximately 10 generations ago, followed by a general recovery.

This pattern is not universally observed in all the American populations: the Brazilian samples from Bambui showed a general decline in population size for the African and European ancestry, according to previous surveys reporting its low heterogeneity ^25^. Conversely, the European ancestry for European Americans (EuroAme) does not show signs of demographic decline, possibly reflecting multiple European waves contributing to this population.

When evaluating the Native American ancestry, the Mexican sample differs from all the others not showing any decrease in the effective population size. A similar behavior was shown when the two samples from Peru were pooled together (Supplementary Figure 10), and could reflect admixture among different Native American groups occurred after the European colonization, or different demographic histories across various American regions.

For the European ancestry, Puerto Ricans (PUR) and Colombians (CLM) showed the most severe decline in effective population size (Figure 3, Supplementary Figure 9).

**Figure 3.**
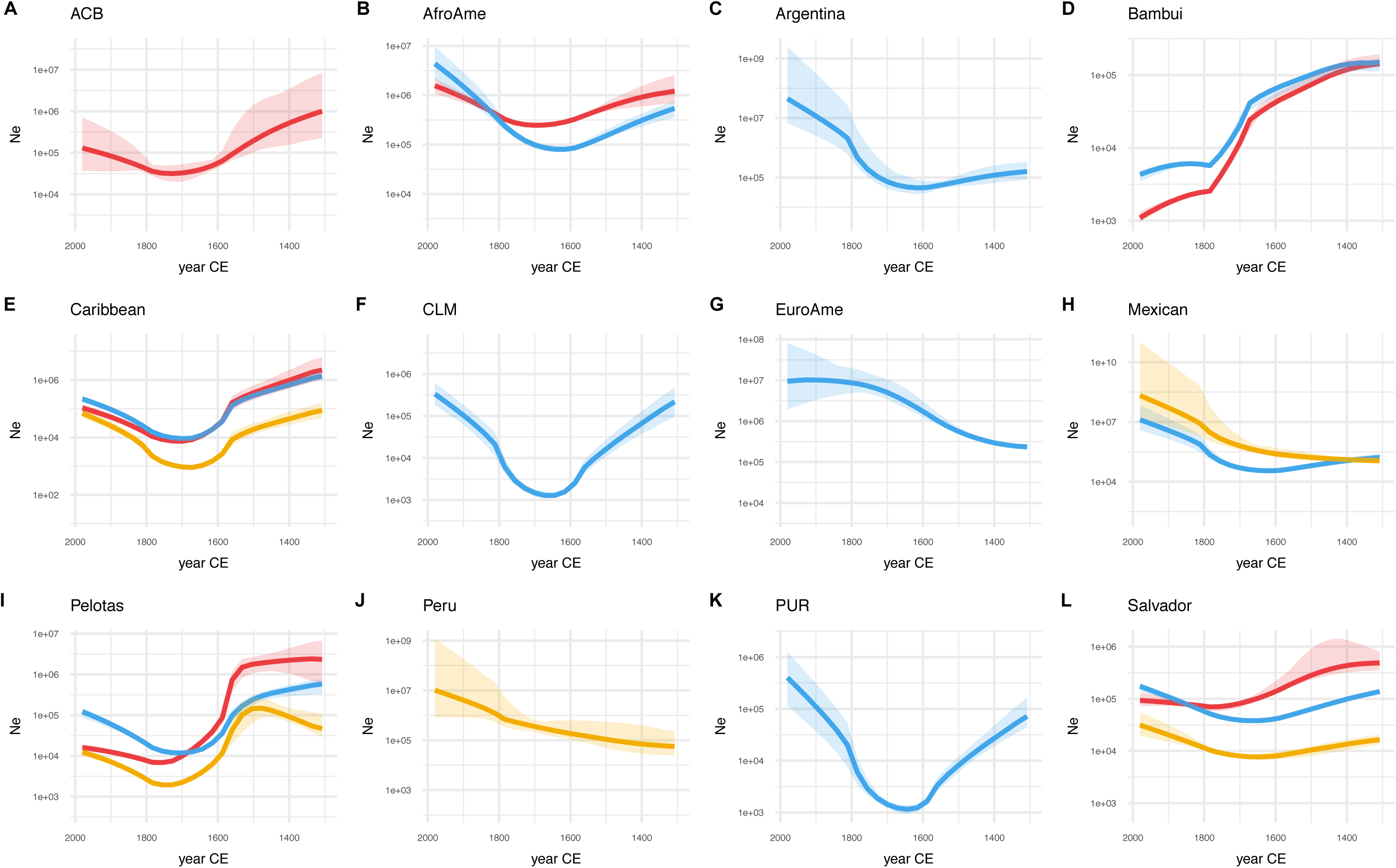
Ancestry-specific effective population size of American populations. We combined Identity by Descent and Local ancestry inferences to estimate ancestry-specific population size through time. The x-axes show time expressed in years of Common Era. The y-axes show ancestry-specific effective population size (Ne), plotted on a log scale. Solid lines show estimated ancestry-specific effective population sizes (red = African ancestry, blue = European ancestry, yellow = Native American ancestry), with ribbons indicating the 95% confidence intervals. Only the population ancestries in which α(continent)* N > 50 where α is the proportion of a specific ancestry and N is the total number of chromosomes in the analysed population are represented.

Interestingly, for the four populations showing a decline-recovery pattern and for which the effective population size for African and European components were available, the African ancestry started to recover later than the European one, with the exception of the Caribbean population. Furthermore, when all the available data points are considered, the time of the last minimum before the recovery is significantly larger for the African ancestry (Wilcoxon test, p-value < 0.05).

## Discussion

Despite being virtually isolated from the rest of the world until five hundred years ago, most of the individuals living in the Americas harbour, together with Native American ancestry, a substantial genomic proportion inherited from Europe and Africa. These ancestral mosaics are the consequence of admixture events occurred after European exploration and colonization, which was followed by African deportation and labour migration that have impacted the American continents in the 19^th^ and 20^th^ century.

The investigation of the times of admixture among the two continents revealed that all the present day American populations are the result of at least one admixture event involving Native American, African, and European sources within the last 6-12 generations, corresponding to 1644 Common Era (CE) and 1812 CE (considering a generation time of 28 years; Figure 1 and Figure 2). However, the approach considering populations does not capture the high complexity of the admixture dynamics, characterized by several waves of migration in the last five centuries, as reported in historical and anthropological records ^7,27,35^. One way to partially overcome this limitation is analysing single individuals rather than populations, capturing a higher degree of variation in the fragment length distribution. Our per-individual time estimations provided several insights into the complexity of admixture in the Americas. It has been recently reported that the origin of Africans deported in the continents followed a general North-South temporal pattern ^36^, with slaves from Senegal and Gambia being deported earlier than the ones from more southern areas (www.slavevoyages.org). In accordance with historical data, the inferred admixture dates involving populations from Senegal and Gambia are older than the ones involving all the others, indeed this area remained the main slave trade site for the Spanish possessions until 1640 ^37^. Similarly, all the dates involving clusters related to Angolan and Namibian individuals are characterized by younger recent admixture times (Figure 2D).

For European sources, estimated admixture dates involving gene flow from Iberia are older than dates of admixture from France/Great Britain sources, which, in turn, are older than admixture events from Italian sources, that, according to historical records became substantial only in the second half of the 19^th^ century.

Furthermore, we assessed the large impact of the Atlantic Slave trade in several populations under study, with patterns reflecting historical records ^27,35,37^.

In detail, our analysis revealed that West-Central Africa ancestry is the most prevalent in the American continents as previously reported ^14,17^, but we additionally identified a high contribution from Senegal and Gambia in Caribbean, Mexico and Colombia in accordance with African slave arrivals predominantly to Spanish-speaking America until 1620s ^27^.

Subsequently, according to disembarkment records, about 50% of all West African slaves were deported to Dutch, French and British sugar plantations in the Caribbean. Accordingly, we estimated a high contribution from Benin and Nigeria in all the Caribbean populations and in populations from US in line with the reported slave arrivals.

Among all the analysed populations, ACB (Barbados) is characterized by the highest Sub-Saharan ancestry proportion (∼88%), possibly due to the presence of sugar cane industry combined with the relatively low European immigration ^35^ in the 18^th^ century.

At a microgeographic scale, Barbadians derives their African ancestry from “*BeninNigeria*” (∼50%) and from “*BeninIvoryCoast*” (∼21%) (Figure 1A and Supplementary Table 3), two of the main source areas reported for the British-mediated slave trade.

In contrast, Brazil shows a peculiar African ancestral composition, characterized by a high proportion of ancestry related to Angola and Namibia, consistent with the Portuguese settlement in Angola from the beginning of the XVII century. A similar African component is also observed in Argentina, probably due to the fact that slaves arrived primarily from Brazil via the Portuguese slave trade from Angola ^36,38^.

The Atlantic coast of Africa was not the only region involved in slave deportation; in the last decades of the slave trade period, Mozambique was the third largest supplier of slaves^27^. We found ancestry related to Southern East African groups in a non-negligible proportion of individuals from Bambui and Pelotas.

While similar works ^14,17^ analyzed Bantu populations from Southern, Southern Eastern and Eastern Africa, here we included Bantu populations from Angola, which has been documented as one of the main regions for slave deportation. Considered together, this study and Gouveia et al. ^17^ suggest an important role of Southwestern, South and Southeastern Africa in shaping the African gene pool of populations from the Atlantic Coast of the Southern Cone of South-America.

For European sources, we confirmed the large impact of Great Britain, France and Iberian Peninsula for all the tested populations, with a distribution reflecting the geographic occupation of the Americas in the Colonial Era.

Furthermore, our approach revealed the existence of several European secondary sources that contributed to many American populations. In fact, we have identified ancestry closely related to Italian populations in European Americans from the United States, Argentinian and Brazilian populations ^39^.

The Italian migration in the Americas has been recently described as one of the largest migrations of the 19^th^ century, and has been usually referred to as the “Italian diaspora” ^40–42^. Although it started soon after 1492, it reached high proportions only in the second half of the 19^th^ century, with more than 11 million individuals migrating towards the continents, largely to the US, Brazil and Argentina.

Between the 1866 and the 1916, approximately 4 million Italians were admitted in the United States. In the 2017 United States Census Bureau nearly 17 million people (5% of global population) were reported as Italian, with proportions spanning from 1.3% to 17% in different states.

In Brazil, also thanks to subsidies offered by the society for the promotion of immigration, after 1820 nearly half of all immigrants were Italians, and in 1876, their annual arrival rate became higher than the one from Portugal. These migrations continued steadily until 1902, when a decree of the Italian government put an end to all subsidized emigration to Brazil ^43^. We found signals of these migrations, mostly related to North Italy, in all the three Brazilian samples analysed, with the highest proportion in Pelotas, followed by Bambui and Salvador.

In Argentina, the identified Italian contribution is related both to the Northern and Southern part of the peninsula, which is in accordance with movements of millions of individuals from Northern (earlier) and Southern (later) Italy registered from the second half of 1800 throughout the 1950s ^9,29^. It has been reported that Italian immigration was the highest (39.4%) compared to the ones from other countries at the beginning of the 20^th^ century ^44,45^.

Therefore, at a pan American level, the distribution of the Italian components is heterogeneous and closely reflects the one reported by historical records.

Moreover, Pelotas is also characterized by contributions from additional sources, such as Central and North-Europe (“*GreatBritain1”*, “*France”*, “*CentralEurope1-2”* and “*Scandinavia”*) in accordance with historical records.

Recently, a survey employing similar methods on five Southern American populations identified South and East Mediterranean ancestries across Americas, which has been interpreted as a contribution from Converso Jews ^24^. Our analysis of the individual ancestry distribution confirmed the presence of Jews and Levantine ancestries in virtually all the analyzed populations, including those from the Caribbean (Supplementary Figure 5).

By evaluating the continental ancestry estimates using an allele frequency method we were able to confirm the sex-biased admixture dynamics suggesting that a higher number of American females than have contributed to the modern populations. Conversely, European males had a larger contribution than females from the same continent.

In contrast, for the African ancestry we observed an inconsistent result, with some, but not all the populations showing evidence for a higher female contribution, partially discordant with historical reports. A possible explanation might be that the ratio between African males and females is lower than the one observed for the European component, preventing its identification with small sample sizes, and suggesting that such patterns (or their absence) should be interpreted with caution, as previously suggested ^33^.

All these results confirm that the European and African components are playing an important role in shaping the genetic differentiation of different American groups, although their demographic evolution after the arrival in the “new world” is still unknown.

The analysis of ancestry-specific effective population sizes demonstrated that, regardless of their composition, virtually all the continental ancestries experienced a general decrease until approximately 10 generations ago, after which a general population size recovery was inferred (Figure 3, Supplementary Figure 9).

Interestingly, the recovery of the African population component postdates those of the European one, possibly reflecting the different conditions experienced by African slaves and European settlers.

On the other hand, the effective population size of the Native American component in Mexicans and Peruvians does not show evidence of decrease, in contrast with historical records reporting a general dramatic decline of the Native American population after European colonization.

This observation is in line with Browning et al. ^34^ in which a smaller reduction in the effective population size of Mexicans for Native American ancestry compared to other populations was observed. This result is also in line with our GLOBETROTTER results, where we found evidence for admixture between two Native American related sources around 15 generations ago.

It may be possible that, the reported decline did not heavily affect the genetic variability of survivor populations; or that individuals from different isolated native groups have been put in contact as a consequence of the European colonization and deportation, as recently suggested for Peruvian populations ^46^. This would result in an inflated effective population size estimate, as we observe in our IBDNe analysis.

In conclusion, we demonstrated that the European and African genomic ancestries in American populations are composed of several different sources that arrived in the Americas in the last six centuries, dramatically affecting their demography and mirroring historical events. The analysis of high quality genomes from the American continents, combined with the analysis of ancient DNA and denser sampling will be crucial to better clarify the genetic impact of these dramatic events. In addition, the fine scale composition here reported is important for the future development of epidemiological, translational and medical studies.

## Methods

### Dataset

We assembled ^11,15,16,20,21,23,25,32,47–69^ a genome-wide dataset of 25,732 worldwide individuals genotyped with different Illumina platforms. Of these, 25,455 were retrieved from publicly available and controlled access resources. In order to increase our resolution in identifying the source of analysed individuals, we added 277 samples from 35 Eurasian populations. Genotype data for 89 samples are available at http://evolbio.ut.ee/. The remaining samples will be available in dedicated future publications.

The obtained dataset was filtered using PLINK ver. 1.9 ^70^ to include only SNPs and individuals with genotyping success rate > 97%, retaining a total of 251,548 autosomal markers.

We used KING to remove one random individual from pairs with kinship parameter higher than 0.0884 ^71^. The final dataset was therefore composed of 17,722 individuals from 261 populations ^11,15,16,20,21,23,25,32,47–69^ (Supplementary Table 1A-B, Supplementary Figure 1). Of these, 11,607 individuals belonging to 22 admixed American populations were treated as ‘recipients’, while the remaining 6,115 samples from 239 source populations were considered ‘donors’.

### PCA analysis

Principal Components Analysis (PCA) was performed on the final dataset using the command --pca from PLINK 1.9. The resulting plot is shown in Supplementary Figure 11.

### Phasing

Germline phase was inferred using the Segmented Haplotype Estimation and Imputation tool (ShapeIT2) software ^72^, using the HapMap37 human genome build 37 recombination map.

### Clustering of donor populations

As a first step, we clustered the individuals belonging to ‘donor’ populations into homogenous groups. First, we used the inferential algorithm implemented in CHROMOPAINTER (v2) ^18^ to reconstruct each individual’s chromosomes as a series of genomic fragments inherited (copied) from a set of donor individuals, using the information on the allelic state of recipient and donors at each available position. Briefly, we ‘painted’ the genomic profile of each donor as the combination of fragments received from other donor individuals. We used a value of 288.998 for the nuisance parameters ‘recombination scaling constant’ (which controls the average switch rate of the HMM) Ne, and 0.00076 for the ‘per site mutation rate’ M, nuisance parameters, as estimated by 10 iterations of the expectation-maximization algorithm in CHROMOPAINTER. This algorithm finds the local optimum values of these parameters iterating over the data. Given the computational complexity of this process, the estimation of these two parameters was obtained by averaging the values calculated from an analysis performed on a subset of six hundred individuals from all the analysed populations, with sample sizes mirroring the global composition of the dataset for five randomly selected chromosomes (3, 7, 10, 18 and 22).

Second, we analysed the painted dataset using fineSTRUCTURE^18^, in order to identify homogeneous clusters. We ran the software in three subsequent steps: the first, also called “greedy”, infers in a fast way a rough clustering summarizing the relationships among individuals, and it is usually used when the number of samples is large (> 5000 individuals); the second, starting from the greedy clustering, performs 1 million MCMC iterations thinned every 10,000 and preceded by 100,000 burn in iterations. This generated a MCMC file (.xml) that was used, by the third run, to build the tree structure using the option --T 1 ^73^.

FineSTRUCTURE classified the analysed individuals into 370 clusters (Supplementary Figure 12). In order to increase the interpretability of subsequent analysis we reduced the number of identified groups. In doing so, we iteratively climbed the tree, and lumped pairs of clusters until the minimum pairwise Total Variation Distance (TVD) estimated on the chunkcounts was lower than a given threshold. Taking into consideration the within continents variability and their relevance as sources to American populations, we applied a threshold of 0.04 for Sub-Saharan African, Asian and Oceanian clusters, 0.03 for North-African, Native American and North-East European clusters and 0.015 for Central, West and South European clusters. After refining, 89 clusters remained (Supplementary Table 2, Supplementary Figure 2). One cluster composed of less than five individuals was excluded from the following further analysis.

### Painting of the recipient populations

We used CHROMOPAINTER, to paint each recipient individual as a combination of genomic fragments inherited by ‘donor individuals’ pooled using the clustering affiliation obtained as previously described, and with the same nuisance parameters inferred for the donor individuals.

### Bayesian haplotype-based ancestry estimation (SOURCEFIND)

We applied a recently developed Bayesian method, SOURCEFIND, ^24^ to estimate the ancestral composition of recipient individuals. Thus, we modelled the copying vector (obtained with CHROMOPAINTER analysis) of each admixed individual as a weighted mixture of copying vectors from the donors. We used as parameters: self.copy.ind=0, number of total (num.surrogates) and expected (exp.num.surrogates) surrogates equal to 8 and 4 respectively; performing (total number of MCMC iterations) 200,000 iterations thinned every 1,000, and preceded by a burn in step of 50,000. Furthermore, we assigned equally-sized proportions to the surrogates (num.slots=100). For each recipient individual, we combined 10 independent runs extracting and averaging the estimates with the highest posterior probability, weighted by their posterior probability. The efficacy and reliability of the method has been assessed for a similar scenario through an extensive simulation approach in Chacón-Duque et al. ^24^.

### Non-Negative Least Square haplotype-based ancestry estimation

CHROMOPAINTER provides a summary of the amount of DNA copied from each donor population. We identified the most closely ancestrally related donor population for each admixed population by comparing their copying vectors to copying vectors inferred in the same way for each of the donor clusters, using a slight modification of non-negative least square (NNLS) function in R 3.5.1 ^74^, and following the approach reported in Montinaro et al. and Leslie et al. ^14,73^. Briefly, this approach identifies copying vectors of donor populations that better match the copying vector of recipient populations as estimated by CHROMOPAINTER. For each recipient population, we decomposed the ancestry of that group as a mixture (with proportions summing to 1) of each sampled potential donor cluster, by comparing the ‘copying vector’ of donor and recipient populations.

### Estimation of admixture dates

In order to provide a temporal characterization of the admixture events in the Americas, we estimated times and most closely related putative sources using population-based and individual-based painting profiles.

In the “population” approach, given the high demand of computational resources requested for the analysis, we have used fastGLOBETROTTER, which, based on GLOBETROTTER ^75^, implements several optimizations in performance, making it suitable for large datasets. In detail, we first harnessed the painting profiles obtained by CHROMOPAINTER by testing for any evidence of admixture using the options null.ind=1, prop.ind=1, and performing 100 bootstrap iterations. For each of the admixture events inferred, we considered only those characterized by bootstrap values for time of admixture between 1 and 400. Subsequently, we estimated time of admixture repeating the same procedure with options null.ind=0 and prop.ind=1.

For the individual analysis we estimated admixture times with GLOBETROTTER, applying the prop.ind=1, null.ind=0 approach to the 11,607 target individuals. In order to remove individuals with “unusual” painting profiles, only those falling in the 2.5-97.5% admixture time confidence interval were retained.

We tested significant differences in times of admixture involving specific African or European clusters by applying a Wilcoxon test using R and setting alternative to “greater”.

### Ancestry-specific effective population size estimation

In order to estimate ancestry-specific effective population size for the 22 recipient American populations we followed the pipeline presented by Browning et al. ^34^(http://faculty.washington.edu/sguy/asibdne/posted_commands.txt).

We used IBD and LA inferred from genome-wide data as a first step. We inferred IBD segments using the refined IBD algorithm implemented in Beagle 4.1, with the following parameters: ibdcm=2, window=400, overlap=24 and ibdtrim=12, as suggested in Browning et al.^34^. Subsequently, we ran the merge-ibd-segments.12Jul18.a0b.jar script to remove breaks and short gaps in the inferred IBD segments (gaps shorter than 0.6cM).

We estimated the local ancestry for genomic fragments in the American individuals using RFMIX. As reference populations we used Yoruba (YRI), Gambia (GWDwg) and Mozambique for Africa, Chinese Han (CHB) and Japanese (JPT) for Asia, Spanish (IBS), British (GBR) and Tuscany (TSI) for Europe and Tepehuano, Wichi and Karitiana for Native American ancestry. We used “PopPhased”, “-n 5” and “--forward-backward” options as recommended in RFMix manual. Then, we corrected the initial phasing following the modifications of RFMIX and using the rephasevit.py script provided by Browning et al. ^34^.

We combined the results from IBD analysis and LA assigning to each IBD segment the most probable ancestry.

Subsequently, we calculated the adjusted number of pairs of haplotypes for each ancestry. This is required because two haplotypes can only be in IBD with respect to a given ancestry at genomic positions if both haplotypes have that ancestry. Therefore, in a sample composed by *n* individuals the ancestry-adjusted number of pairs of haplotypes is equal to:

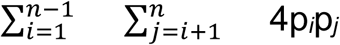

(where *i* and *j* are independent individuals and *pi* and *pj* are their proportions of the given ancestry).

Finally, we used the obtained “npairs” to run IBDNe software (version ibdne.07May18.6a4) ^34,76^ in default mode, except for filtersample=false.

### Sex-biased admixture evaluation

We intersected SNPs from the X chromosome that were present in both our main datasets and in the 1000 Genomes Project samples. Three admixed American groups (Mexican, Maya and Mayas), were removed because the data did not include any genotypes for chromosome X. We revised and imputed sex assignments based on X chromosome data using the --impute-sex command in PLINK. A male or female call is made when the rate of homozygosity is >80% and <20%, respectively. Individuals for which the sex imputation was ambiguous were removed and heterozygous SNPs in male X chromosomes were set as missing. After this step, only samples and positions with a genotyping rate >= 97% were retained: 5,227 SNPs in a total of 15,353 individuals. The same set of individuals was extracted from the filtered autosomal dataset with 258,720 SNPs. Subsequently, we performed LD pruning (--indep-pairwise 200 50 0.2) in both X chromosome and autosomal data sets, resulting in a total of 2,519 and 116,912 SNPs, respectively. We ran separate unsupervised ADMIXTURE (version 1.3.0^30^) analysis for the two datasets using K values=3 and 10 independent runs. We used the option ‘--haploid=‘male:23’ in order to properly treat male individuals and chose the best run according to the highest value of log likelihood. Finally, we performed paired Wilcoxon tests in order to test for significant differences between the ancestry proportions observed in the autosomes versus the X chromosome and used Bonferroni correction for multiple-testing (adjusted p-value < 0.05).

## Supplementary Tables Captions

**S1. Details of the genotype data used in this study at individual (A) and population (B) level.** “D/R” column refers to whether the sample was used as “Donor” or “Recipient” in the CHROMOPAINTER/fineSTRUCTURE pipeline.

**S2. Cluster composition for the 89 clusters obtained after filtering.** We iteratively climbed the tree at each node, and lumped pairs of clusters until the minimum pairwise Total Variation Distance estimated on the chunkcounts was lower than a threshold. Taking into consideration the continental variability and the relevance for their contribution in the Americas, we applied a threshold of 0.04 for Sub-Saharan African, Asian and Oceanian, 0.03 for North-African, Native American and North-East European and 0.015 for Central, West and South European clusters.

**S3. Results of SOURCEFIND analysis.** The table reports the ancestral proportions (as percentages) inferred by SOURCEFIND for the 21 most representative fineSTRUCTURE clusters (contribution ≥ 2% in at least one recipient population). The column “others” contains the sum of the proportions for all the cluster contributing less than 2%.

**S4. Results of Autosomes Vs X admixture analysis.** For each of the 19 populations analysed (Maya, Mayas and Mexican were filtered out) are reported: the number of the samples, the resulting p-values of the performed Wilcoxon tests and the median value of the logarithmic Autosomes/X chromosome ratio.

**S5. Results of GLOBETROTTER considering populations.** For each target population the following parameters are reported: most supported event (“GT- response”), time of admixture event 1 (“time-event1”), proportion of admixture for the minor population for the first admixture event (“alpha-event1”), closer representative for minor contributing population in event 1 (“sourceA-event1”), closer representative for major contributing population in event 1 (“sourceB-event1”), time of admixture event 2 (“time-event2”), proportion of admixture for the minor population for the second admixture event (“alpha-event2”), closer representative for minor contributing population in event 2 (“sourceA-event1”), closer representative for major contributing population in event 2 (“sourceB-event2”), minimum, maximum, 2.5% and 97.5% values for date distribution for both the inferred events (“time1-min”, “time1-max”, “time1-2.5%”, “time1-97.5%”, “time2-min”, “time2-max”, “time2-2.5%”, “time2-97.5%”), p-value for event 1 and event 2 (“p-value1” and “p-value2”, respectively).

## ACKNOWLEDGMENTS

We thank Estonian Genome Center and Andres Metspalu for access to the Georgians, Germans, Hungarians, Latvians, Lithuanians and Ukrainians samples. We thank Doron Behar for early access to the Ashkenazi_Jews, Karaite_Jews, Lemba_Jews data. We acknowledge Elza Khusnutdinova for early access to Avars, Bessermans, Cirkassian, Kabardin, Karelian, Komis, Mordovians, Udmurts and Vepsas data. We thank Rusudan Khukhunaishvili and Sophiko Tskvitinidze for early access to Adjarians data. We thank Oleg Balanovsky for the access to Cossack, Tatars-Crimea-Coast, Tatars-Crimea-Mountain, and Tatars-Crimea-Steppe data. All these samples will be published in dedicated papers. We thank Lena Khushniarevich for Tatars_from_Belarus samples and Vladimir Ferak for some of the Roma samples.

This study received support from the Italian Ministry of Education, University and Research (MIUR): Dipartimenti di Eccellenza Program (2018–2022) - Dept. of Biology and Biotechnology “L. Spallanzani”, University of Pavia (to A.A, A.O., O.S, A.T); the Fondazione Cariplo (project 2018-2045 to A.A, A.O., A.T)). This research was supported by the European Union through the European Regional Development Fund (Project No. 2014-2020.4.01.16-0030 to L.O., M.M., F.M.; Project No. 2014-2020.4.01.16-0271 to R.F.; Project No. 2014-2020.4.01.16-0125 to R.F.; Project No. 2014-2020.4.01.16-0024 to D.M., L.P.)”. This work was supported by the Estonian Research Council grant PUT (PRG243) (to R.F., M.M., L.P.). This work was supported by institutional research funding IUT (IUT24-1) of the Estonian Ministry of Education and Research (to T.K.). This research was supported by the European Union through Horizon 2020 grant no. 810645 (to M.M.). Computational analyses were performed at the High Performance Computing Center of University of Tartu; we thank Tuuli Reisberg for assistance in data management.

## Authors’ contributions

CC, LP, MM and FM initially conceived the idea. FM, ETS, LP and MM developed the plan. LO, FM, LP and RJ performed the analyses. RJK, AK, OB, EB, LA, ST, SM, DMn, LK, AL, NM and CAM shared unpublished data. LO, FM, LP, RF, DMj, MOS and ETS (with all?) wrote the manuscript. All the other authors interpreted results, revised and approved the final version of the manuscript.

## Notes

#### Summary of Updates

Supplemental files updated

