## Supplementary Material for "The genomic impact of European colonization of the Americas"

### Supplementary Material
### "The genomic impact of European colonization of the Americas"

Linda Ongaro[1], Marilia O. Scliar[2,3], Rodrigo Flores[1], Alessandro Raveane[4], Davide Marnetto[1], Stefania Sarno[5], Guido A. Gnecchi-Ruscone[5,6], Marta E. Alarcon-Riquelme[7], Etienne Patin[8], Pongsakorn Wangkumhang[9], Garrett Hellenthal[9], Miguel Gonzalez-Santos[10], Roy J. King[11], Anastasia Kouvatsi[12], Oleg Balanovsky[13], Elena Balanovska[13], Lubov Atramentova[14], Shahlo Turdikulova[15], Sarabjit Mastana[16], Damir Marjanovic[17,18], Lejla Kovacevic[19], Andreja Leskovac[20], Maria F. Lima-Costa[21], Alexandre C. Pereira[22], Mauricio L. Barreto[23,24], Bernardo L. Horta[25], Nédio Mabunda[26], Celia A. May[27], Andrés Moreno-Estrada[28], Alessandro Achilli[4], Anna Olivieri[4], Ornella Semino[4], Kristiina Tambets[1], Toomas Kivisild[29], Donata Luiselli[30], Antonio Torroni[4], Cristian Capelli[10], Eduardo Tarazona-Santos[3], Mait Metspalu[1], Luca Pagani[1,31], and Francesco Montinaro[1,10]

[1]Estonian Biocentre, Institute of Genomics, University of Tartu, Tartu, Estonia
[2]Human Genome and Stem Cell Research Center, Biosciences Institute, University of São Paulo, 05508-090, SP, Brazil
[3]Departamento de Genética, Ecologia e Evolução, Instituto de Ciências Biológicas, Universidade Federal de Minas Gerais, Belo Horizonte, MG, 31270-901, Brazil
[4]Department of Biology and Biotechnology "L. Spallanzani", University of Pavia, Pavia, Italy
[5]Department of Biological, Geological and Environmental Sciences, University of Bologna, Bologna, Italy
[6]Department of Archaeogenetics, Max Planck Institute for the Science of Human History, Jena, Germany
[7]GENYO, Centre for Genomics and Oncological Research: Pfizer/ University of Granada/ Andalusian Regional Government, Granada, Spain
[8]Human Evolutionary Genetics Unit, Institut Pasteur, UMR2000, CNRS, Paris 75015, France
[9]Department of Genetics, Evolution and Environment and UCL Genetics Institute, University College London, London WC1E 6BT, UK
[10]Department of Zoology, University of Oxford, Oxford, UK
[11]Department of Psychiatry and Behavioral Sciences, Stanford University School of Medicine, Stanford, CA
[12]Department of Genetics, Development and Molecular Biology, School of Biology, Aristotle University of Thessaloniki, Greece
[13]Vavilov Institute of General Genetics, Research Centre of Medical Genetics, Moscow, Russia
[14]Department of Genetics and Cytology, V.N.Karazin Kharkiv National University, Kharkiv, Ukraine
[15]Laboratory of Genomics, Institute of Bioorganic Chemistry, Academy of Sciences Republic of Uzbekistan, Tashkent, Uzbekistan
[16]School of Sport, Exercise and Health Sciences, Loughborough University, Loughborough, UK
[17]Department of Genetics and Bioengineering, Faculty of Engineering and Information Technologies, International Burch University, 71000 Sarajevo, Bosnia and Herzegovina
[18]Institute for Anthropological Researches, Zagreb, Croatia
[19]Institute for Genetic Engineering and Biotechnology, Sarajevo, Bosnia and Herzegovina
[20]Vinca Institute of Nuclear Sciences, University of Belgrade, Belgrade, Serbia

[21]Instituto de Pesquisa Rene Rachou, Fundação Oswaldo Cruz, Belo Horizonte, MG, 30190-002, Brazil
[22]Instituto do Coração, Universidade de São Paulo, São Paulo, SP, 05403-900, Brazil
[23]Instituto de Saúde Coletiva, Universidade Federal da Bahia, Salvador, BA, 0110-040, Brazil
[24]Center of Data and Knowledge Integration for Health (CIDACS), Fundação Oswaldo Cruz (FIOCRUZ), Salvador, Brazil
[25]Programa de Pós-Graduação em Epidemiologia, Universidade Federal de Pelotas, 464, Pelotas, RS, 96001-970, Brazil
[26]Instituto Nacional de Saúde, Maputo, Mozambique
[27]Department of Genetics and Genome Biology, University of Leicester, Leicester, UK
[28]National Laboratory of Genomics for Biodiversity (LANGEBIO), CINVESTAV, Irapuato, Guanajuato 36821, Mexico
[29]Department of Human Genetics, KU Leuven, Leuven, Belgium
[30]Department of Cultural Heritage, University of Bologna, Ravenna Campus, Italy
[31]Department of Biology, University of Padua, Padua, Italy

### Contents

**List of Supplementary Figures** 4

**1 Supplementary Text** 5

1.1 Historical introduction . . . . . 5

1.2 Description of FineSTRUCTURE clustering results . . . . . 5

**2 Supplementary Figures** 7

**Bibliography** 19

### 1 Supplementary Text

#### 1.1 Historical introduction

The Colonial Era of the Americas started soon after the European discovery of the continent in 1492, when Spanish and British empires started to explore and settle the Southern and Northern region, respectively[3;1]. The Spanish empire first colonized Caribbean Islands, setting the basis for the later colonization of mainland territories. These conquests heavily impacted autochthonous populations living in the area, which were decimated by wars and new pathogens brought by Europeans. It has been estimated that approximately 90% of the Native American population in Mexico perished following the arrival of new colons[7]; while, for example, the Natives from the cold areas in the Andes survived the catastrophe better than elsewhere decreasing by 20-25% in 30 years. On the other hand, population decline in the warmer northern Andes reached levels comparable to that of Mesoamerica[2]. The "Atlantic slave Trade" occurred between the 16th and 19th century, was initiated by Spanish and Portuguese leading to the presence in contemporary American populations of hundreds of millions of people with African ancestry, with the largest proportions in Brazil, the Caribbean, and the United States. It has been estimated that more than 12 million African slaves arrived in the new continent during that period[5]. During the first three centuries, almost all the slaves came from two coastal areas in Africa: the Bight of Benin and West-Central Africa (www.slavevoyages.org). However, with the increasing demand for slaves during 19th century, the Portuguese government permitted free trade for the Brazilian slavers with all the ports of East Africa; in fact Mozambique became the third largest supplier of slaves in that century, ahead of the bight of Biafra and just behind Benin[4].

#### 1.2 Description of FineSTRUCTURE clustering results

FineSTRUCTURE classified the analysed individuals into 370 clusters (Supplementary Figure 12). In order to increase the interpretability of subsequent analysis we reduced the number of identified groups. To do this, we iteratively climbed the tree at each node, and lumped pairs of clusters until the minimum pairwise Total Variation Distance (TVD) estimated on the chunkcounts was lower than a given threshold. Taking into consideration the within continents variability and their relevance as sources to American populations, we applied a threshold of 0.04 for Sub-Saharan African, Asian and Oceanian clusters, 0.03 for North-African, Native American and North-East European clusters and 0.015 for Central, West and South European clusters. After refining, 89 clusters remained (Supplementary Table 2, Supplementary Figure 2). Finally, one cluster composed of less than five individuals was excluded from the following further analysis.

African individuals were classified onto 40 clusters, with a clear split between sub-Saharans, North and Eastern Africans (Supplementary Figure 2). We identified 7 North African clusters including individuals from Morocco, Egypt and the Levant. Individuals from Western and South-Western Africa (sub-Saharan), i.e. from the major slave trading regions, are grouped in 14 clusters. East African individuals are distributed across 10 clusters, while the cluster *"SouthEastAfrica"* includes individuals from Mozambique and Zimbabwe (together with 10 Bantu South Africans). European individuals are differentiated into 36 clusters, mirroring the geographic location of the analysed samples. We identified two Iberian clusters: *"Spain"* that includes mostly Spanish samples (60 Spanish (92,3%), 4 French and 1 Corsican; Supplementary Table 1) and *"SpainPortugal"* which contains 25 Spanish and all the Portuguese samples (10 individuals). Italians are grouped into four groups, reflecting the peculiarity of Sardinian individuals and the genetic differences among the peninsula[6]. Basques form two region-specific genetic groups, one in France and one in Spain. The British samples fall into two different clusters; the vast majority of the samples (74 individuals) cluster together with 17 Welsh and 4 additional individuals from Germany and Sweden, while a smaller subset (24 individuals) forms a homogeneous cluster with Orcadians, possibly reflecting the northernmost nature of the group. Central-North-Eastern Europe is represented by seven clusters containing individuals from multiple countries such as Lithuania, Poland, Belarus, Hungary, Russia, Germany, Austria, Finland and Norway. Four distinct groups including Jewish individuals were identified (Supplementary Figure 2, Supplementary Table 2). Native Americans (American populations characterized by more than 95% of autochthonous ancestry) are grouped into three main clusters, one composed only by Brazilian samples (Karitiana and Surui), one including only Wichi and one composed by several populations (Piapoco from Colombia, Colla from Argentina, Tepehuano, Zapotec and Pima From Mexico). East Asian and Oceania individuals are grouped into five clusters, each exclusively containing individuals from the same population (Supplementary Table 2).

#### 2 Supplementary Figures

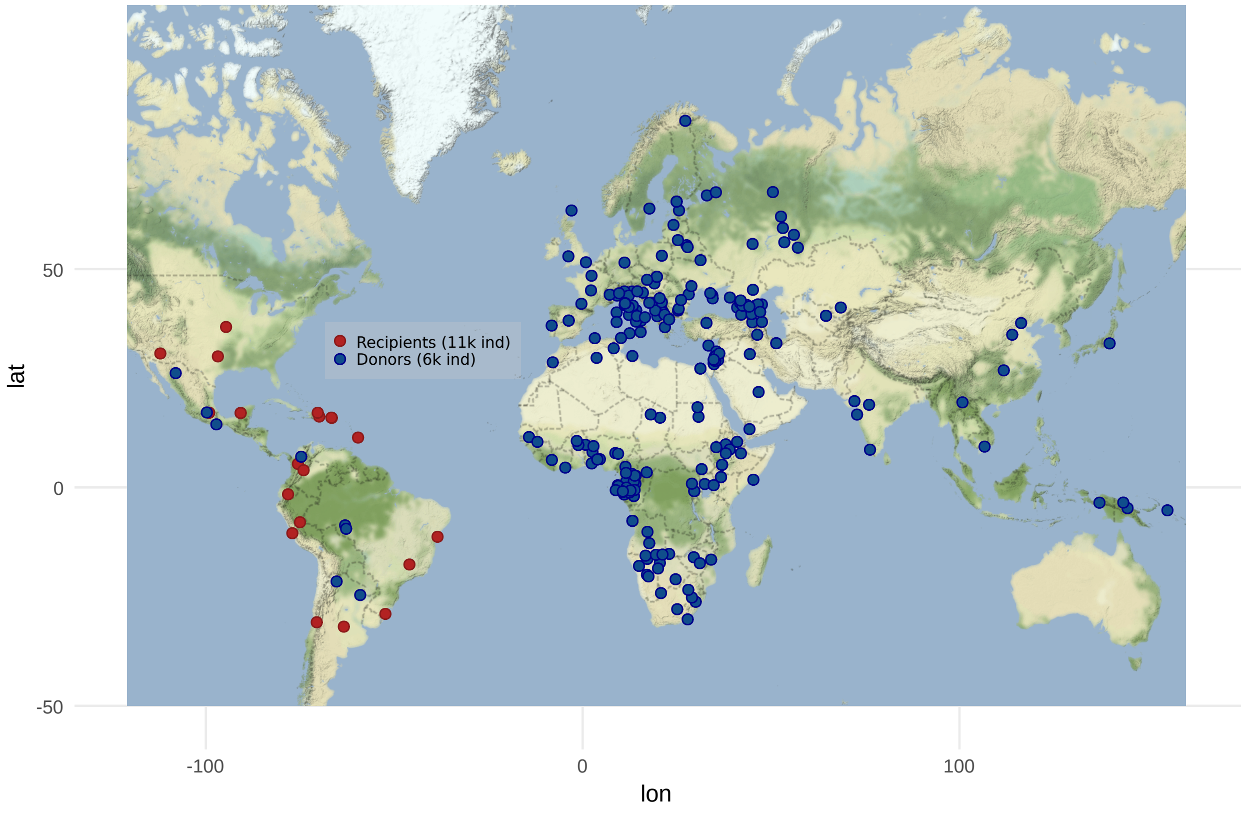

Supplementary Figure 1: Geographic locations of all populations included in the final dataset (see Supplementary Table 1A-B). We analysed 11,607 Recipients and 6115 donors.

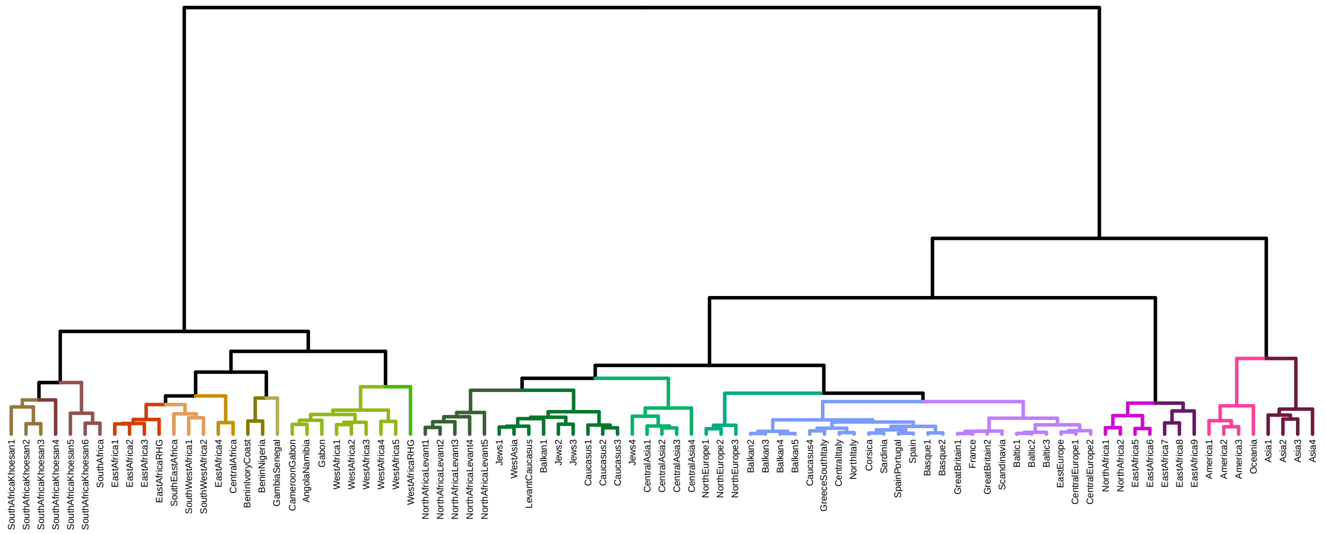

Supplementary Figure 2: Refined fineSTRUCTURE tree of 89 homogeneous clusters (as explained in Methods). Each leaf has been labelled according to the corresponding cluster name reported in Supplementary Table 2, and colored in 20 different macro-groups.

Supplementary Figure 3: Ancestral mosaic of American populations obtained with NNLS analysis. Each barplot shows the genetic composition of admixed populations. Only the contribution for the 26 most representative fineSTRUCTURE clusters (proportion of at least 2% in one recipient population) is reported.

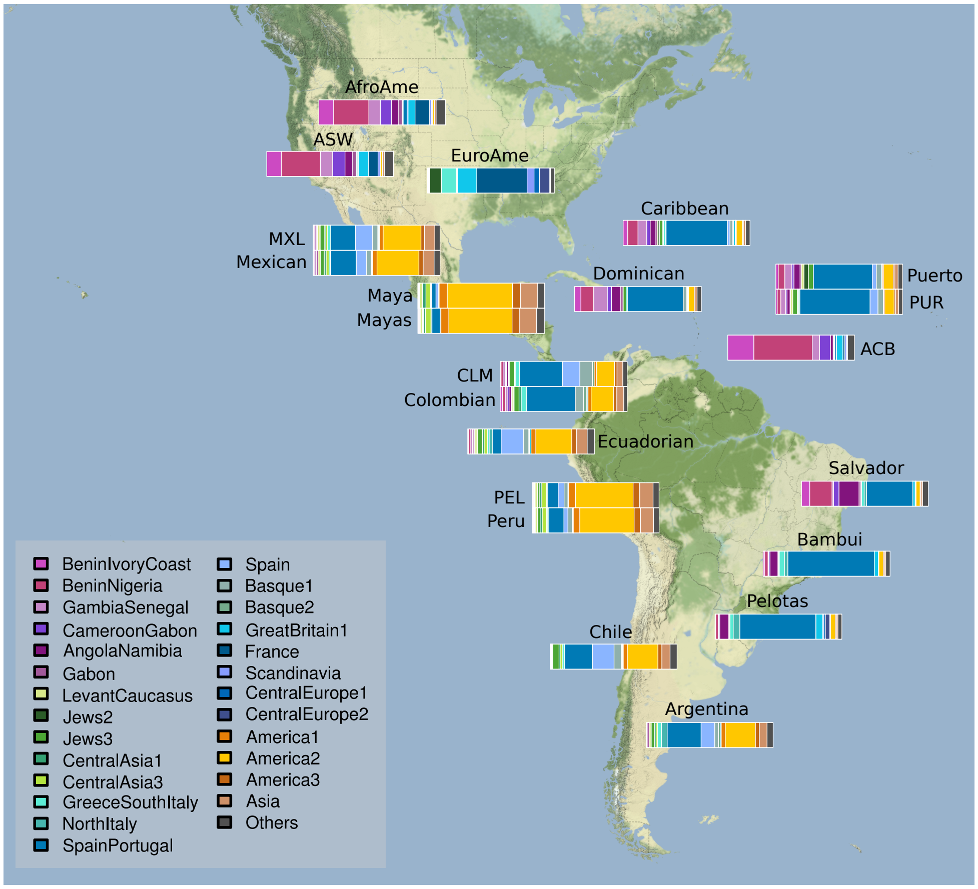

Supplementary Figure 4: The African relative contribution to Americas. We have estimated the relative contribution of the main contributing African sources in Americas. Each barplot shows the proportion of a specific ancestry in different American samples. Only populations having relative ancestries proportions ≥2% are shown.

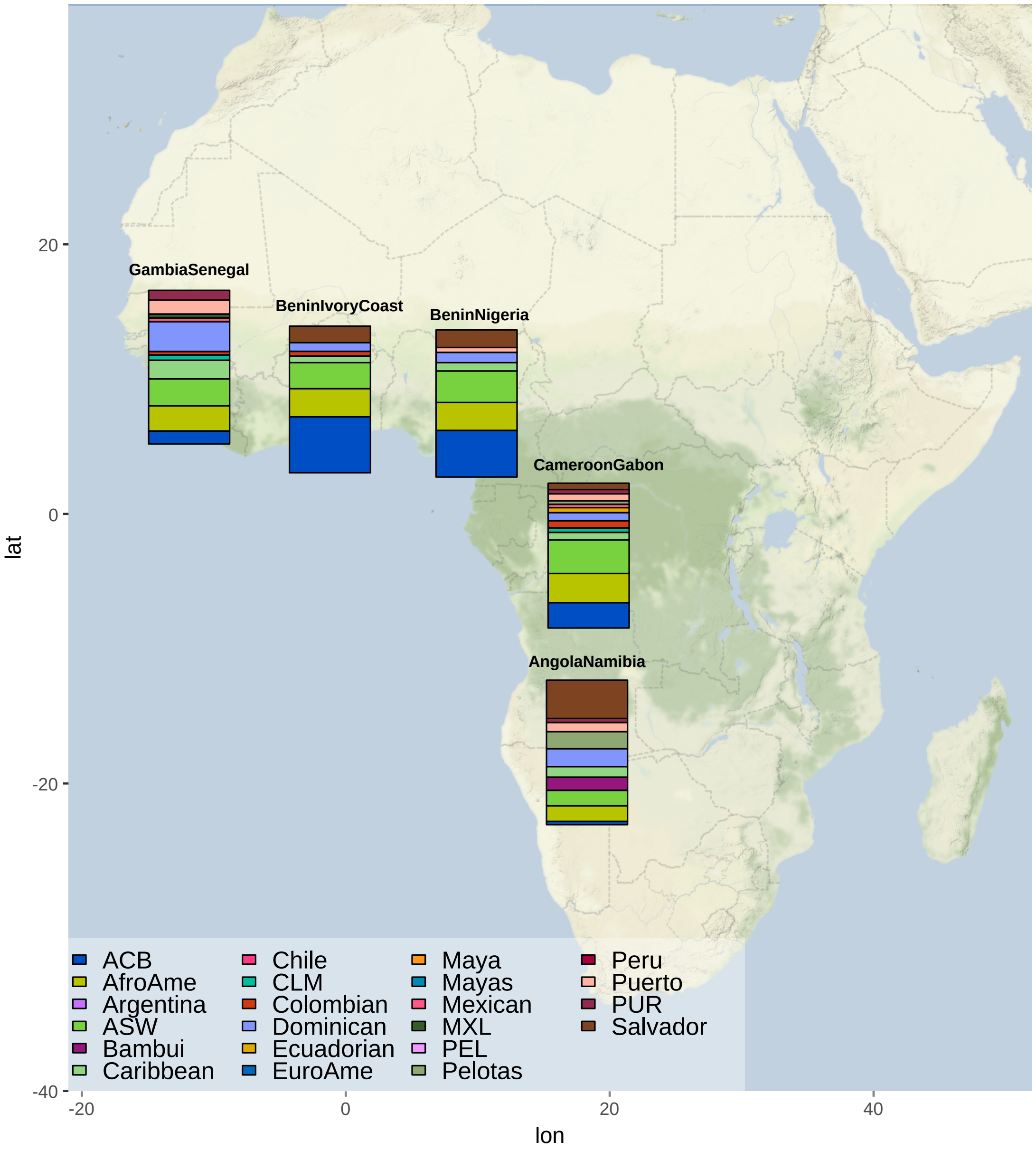

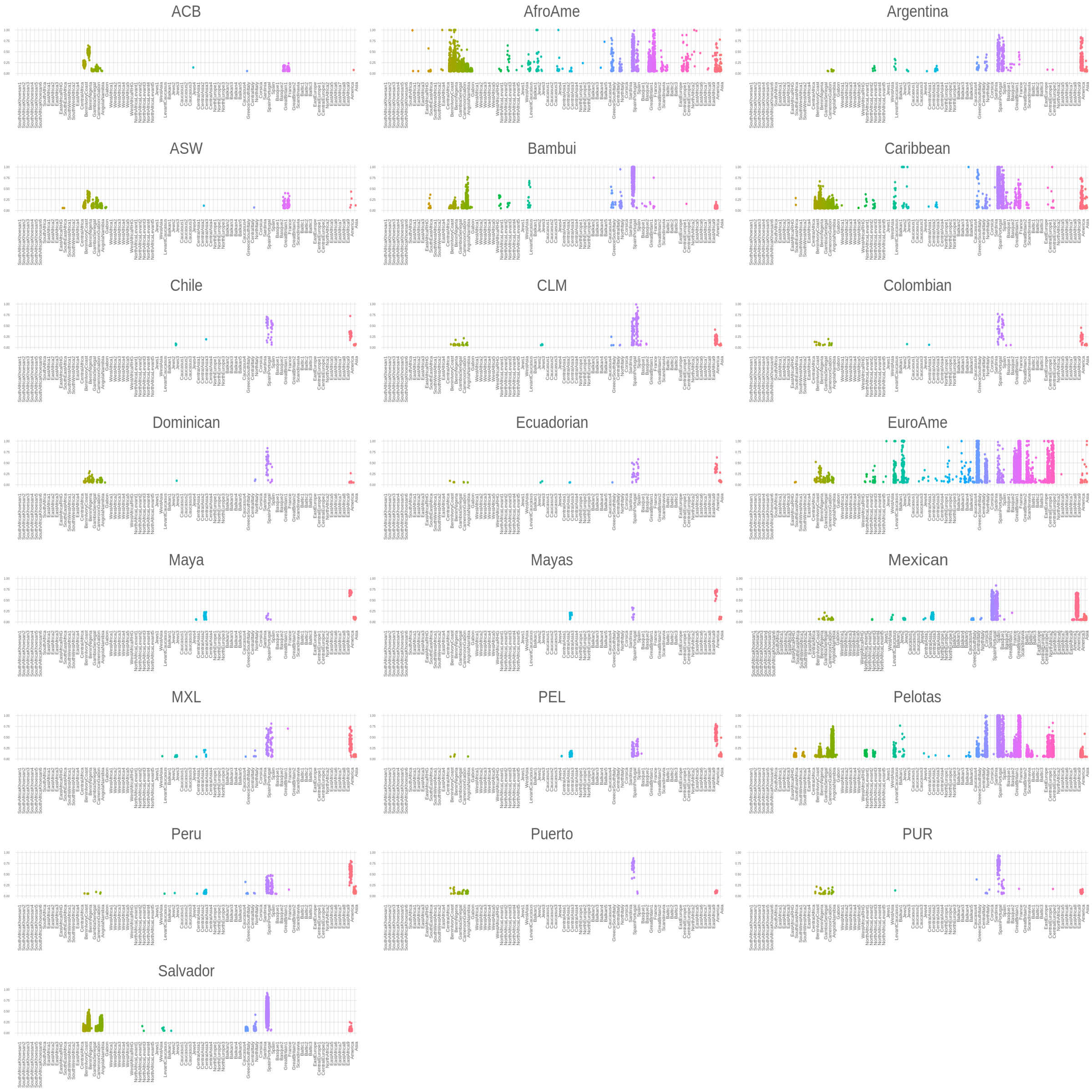

Supplementary Figure 5: Individual ancestral proportion in 11,607 American individuals from 22 populations as inferred by SOURCEFIND. Only points for ancestry proportion >5% are shown.

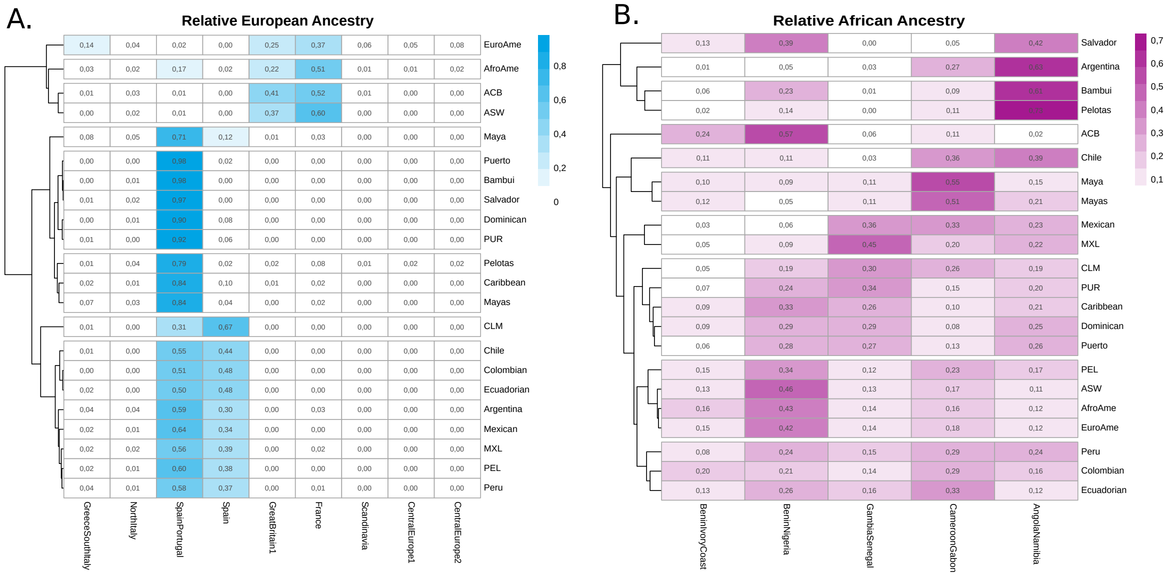

Supplementary Figure 6: Heatmaps and UPGMA dendrogram of European and African relative ancestries. For each population we have considered the relative contribution of each donor cluster into the total European (A) or African (B) ancestry.

Supplementary Figure 7: The European relative contribution to Americas. We have estimated the relative contribution of the main contributing European sources in Americas. Each barplot shows the proportion of a specific ancestry in different American samples. Only populations having relative ancestries proportions $\geq$2% are shown.

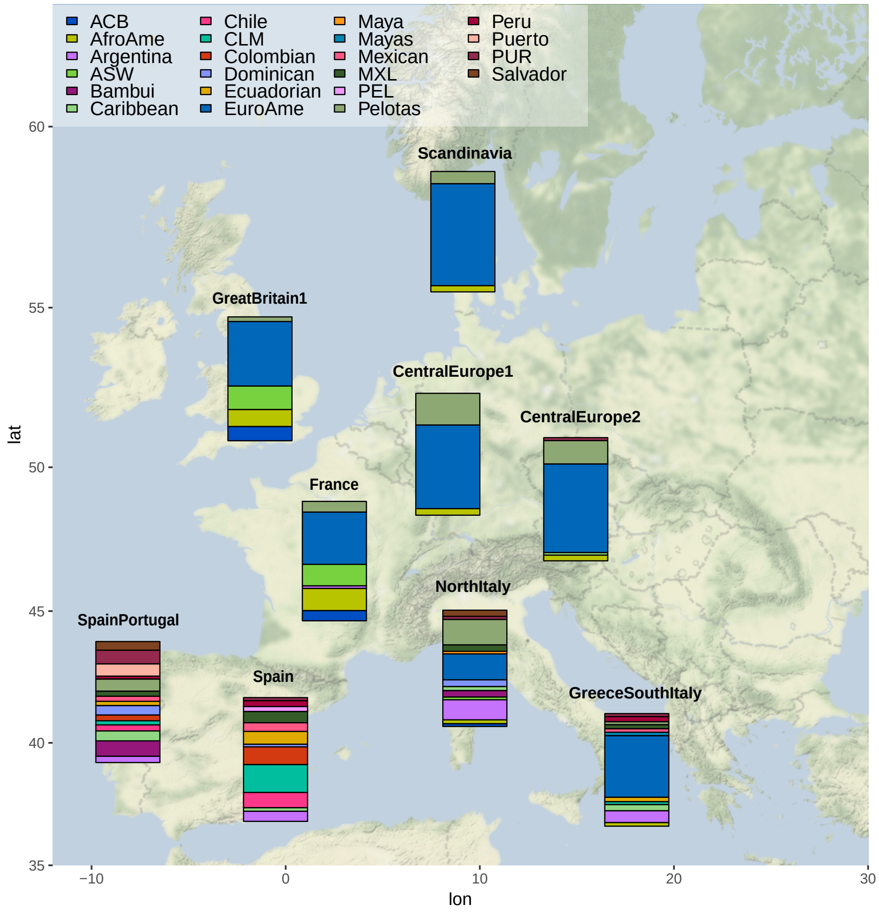

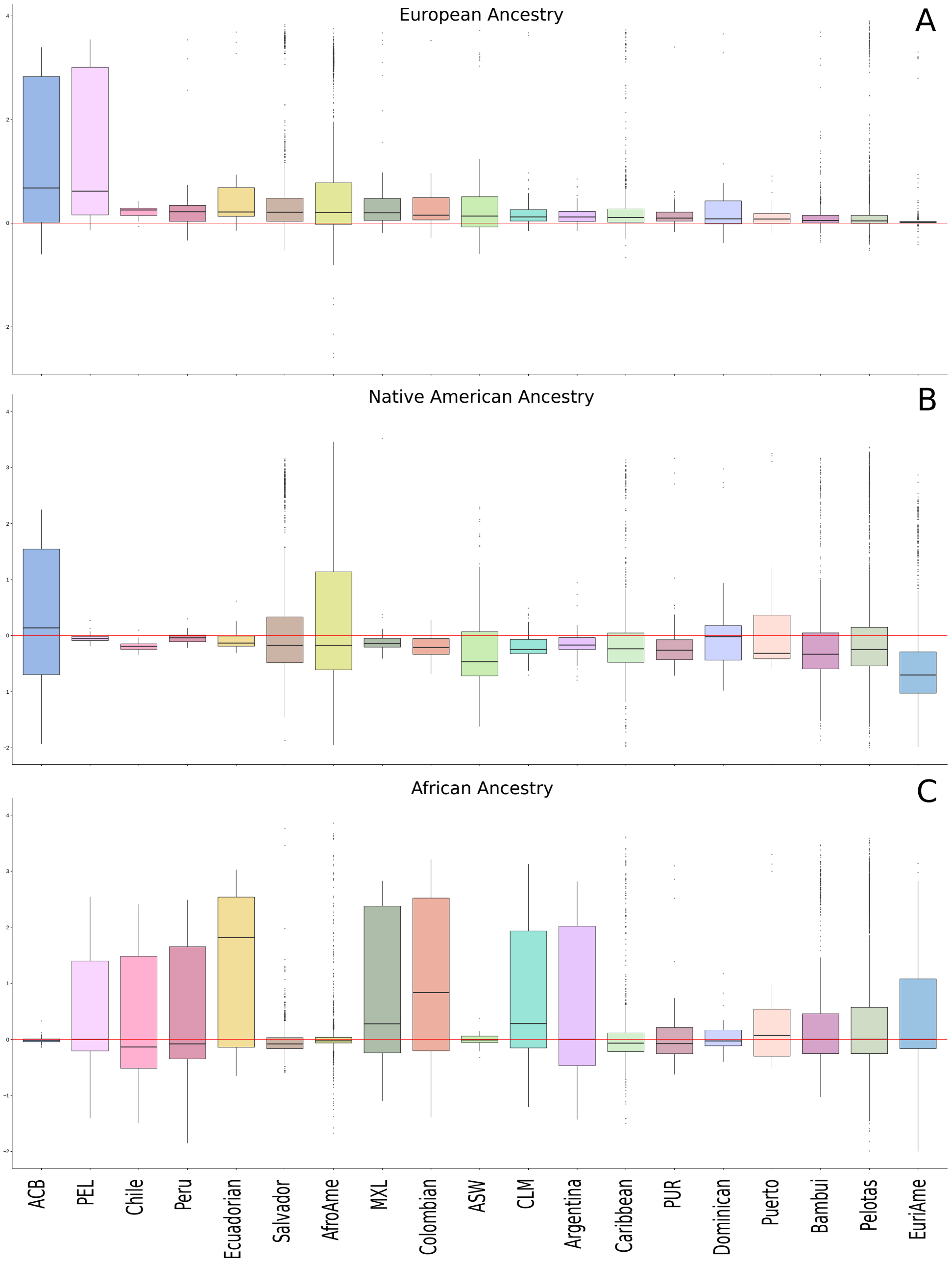

Supplementary Figure 8: Autosomes vs X chromosome ancestry proportions. Each boxplot shows the log10-scaled ratio of autosomal to X chromosome ancestry proportion for A) European, B) Native American and C) African continental components as inferred by ADMIXTURE analysis (K=3) in 19 American populations.

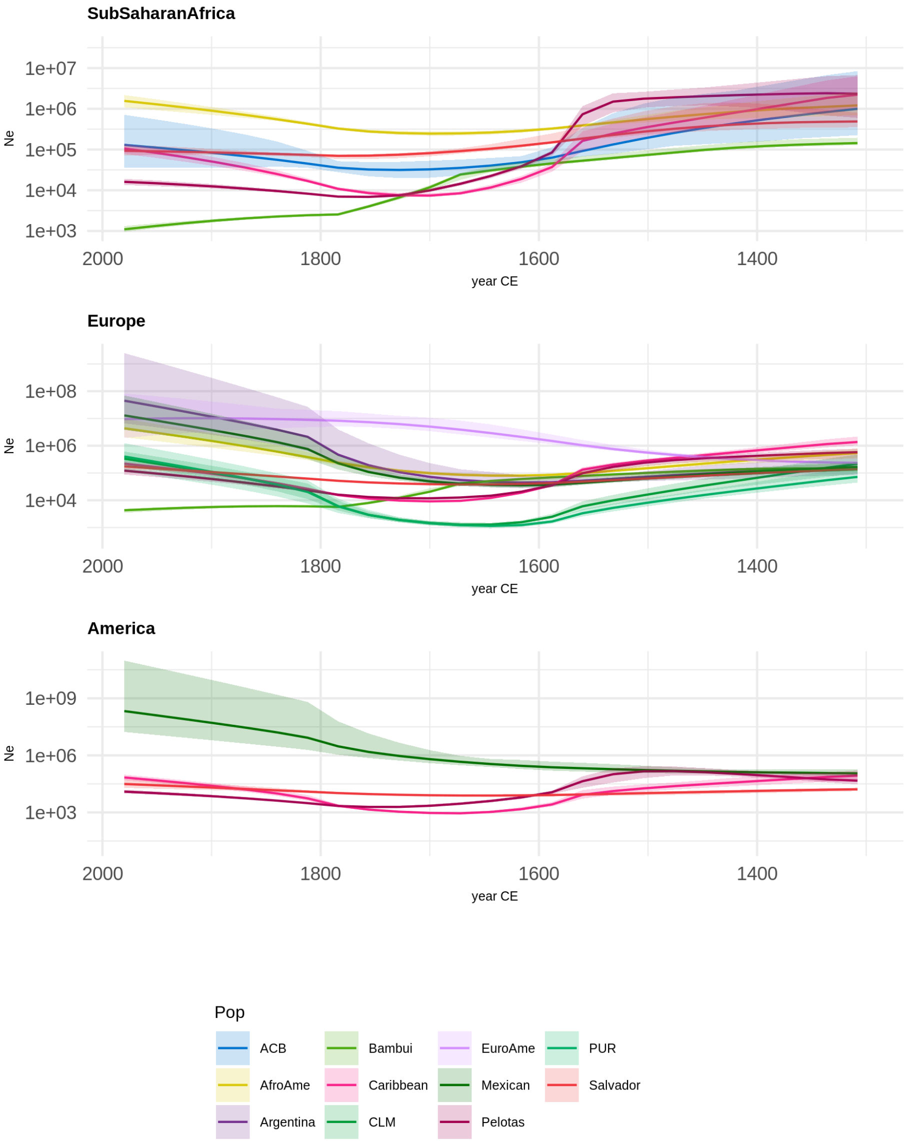

Supplementary Figure 9: Ancestry-specific effective population size of American populations decomposed for single ancestry. We combined Identity by Descent and Local Ancestry analysis inference to estimate Ancestry Specific Population size. The x-axes show time expressed in years of Common Era. The y-axes show ancestry-specific effective population size (Ne), plotted on a log scale. The lines show estimated ancestry-specific effective population sizes, and ribbons indicate the 95% confidence intervals. Only the population ancestries in which α(continent)* N > 50 where αis the proportion of a specific ancestry and N is the total number of chromosomes in the analysed population are represented.

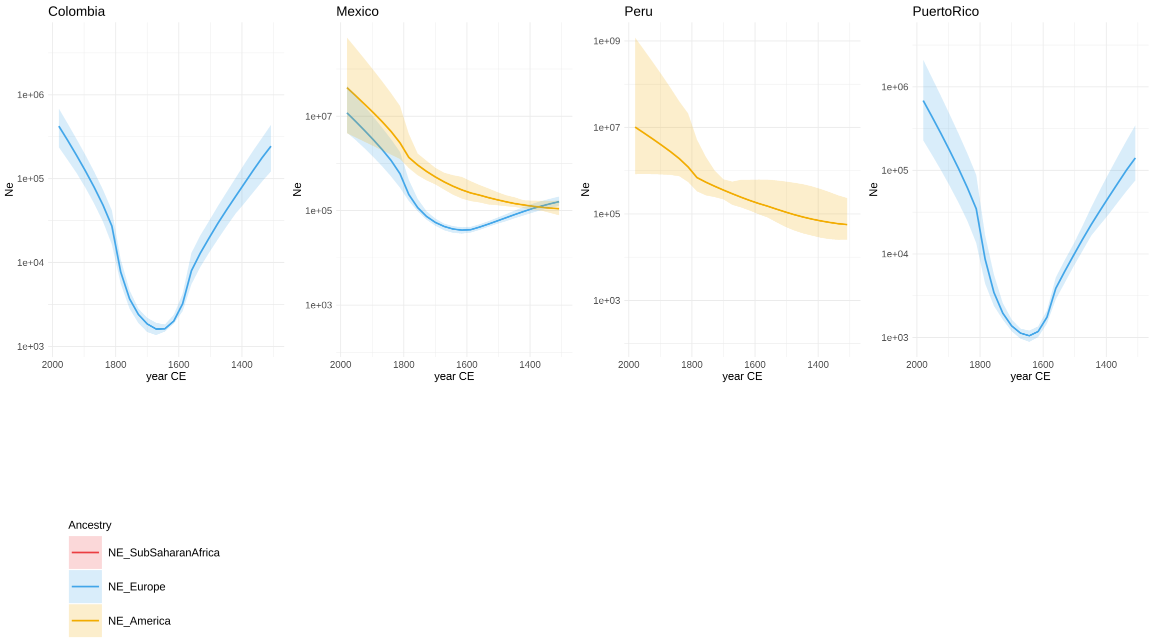

Supplementary Figure 10: Ancestry-specific effective population size of American populations decomposed for samples from the same population. We combined samples from Colombia, Peru, Mexico and Puerto Rico and inferred the Ancestry Specific Effective Population using IBDNe.

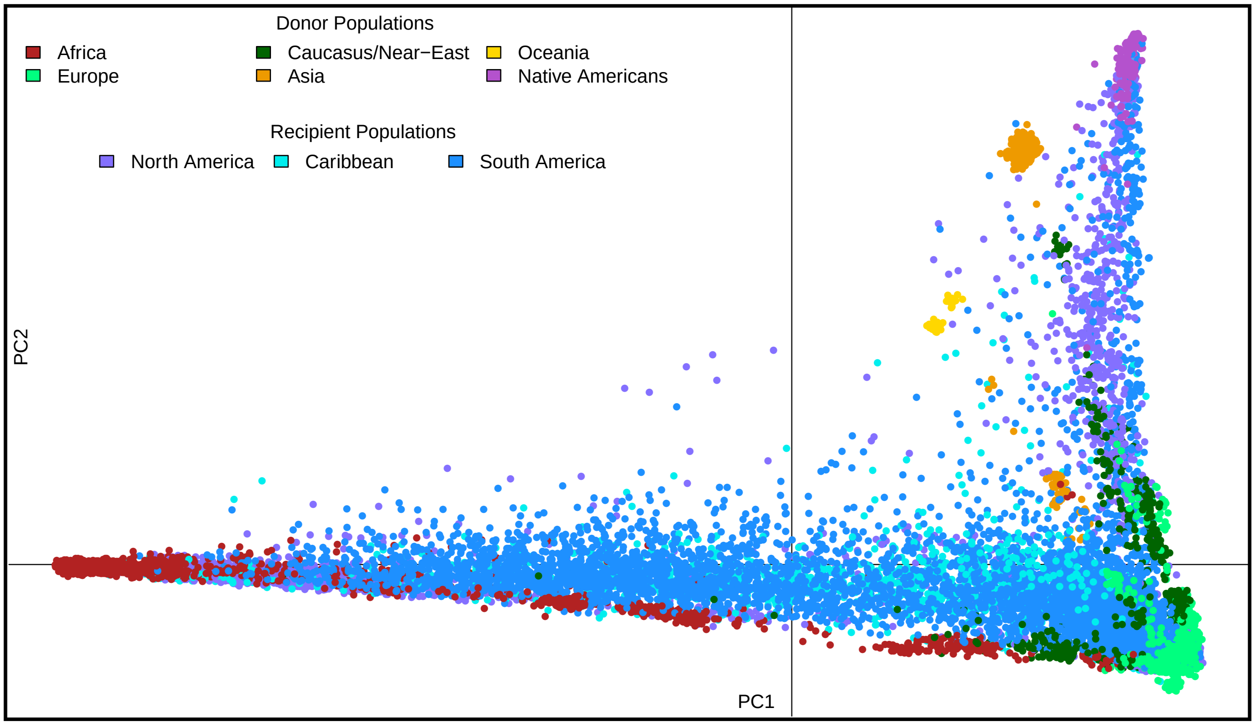

Supplementary Figure 11: Principal Component Analysis for the analysed data.

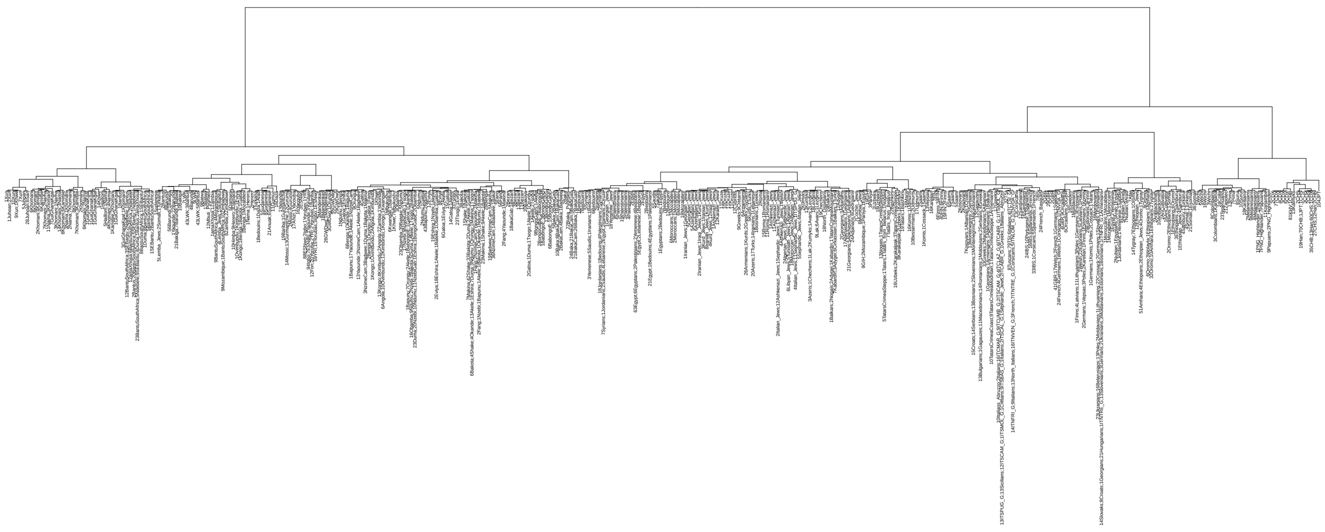

Supplementary Figure 12: Dendrogram of donor individuals clustered by fineSTRUCTURE into 370 clusters. Each branch label is in the form X¡popA¿;Y¡popB¿, where X and Y are the number of individual for popA and popB in the cluster.

[2] Leslie Bethell. *The Cambridge History of Latin America.* Cambridge University Press, December 1984.

[3] Henry Kamen. *Spain's road to empire: The making of a world power, 1492-1763.* Penguin UK, 2003.

[4] Herbert S Klein. *The Atlantic Slave Trade.* Cambridge University Press, April 1999.

[5] Paul E Lovejoy. The impact of the atlantic slave trade on africa: A review of the literature1. *J. Afr. Hist.*, 30(3):365–394, November 1989.

[6] Alessandro Raveane, Serena Aneli, Francesco Montinaro, Georgios Athanasiadis, Simona Barlera, Giovanni Birolo, Giorgio Boncoraglio, Anna Maria Di Blasio, Cornelia Di Gaetano, Luca Pagani, Silvia Parolo, Peristera Paschou, Alberto Piazza, George Stamatoyannopoulos, Andrea Angius, Nicolas Brucato, Francesco Cucca, Garrett Hellenthal, Antonella Mulas, Marine Peyret-Guzzon, Madzia Zoledziewska, Abdellatif Baali, Clare Bycroft, Mohammed Cherkaoui, Christian Dina, Jean-Michel Dugoujon, Pilar Galan, Joanna Giemza, Toomas Kivisild, Mohammed Melhaoui, Mait Metspalu, Simon Myers, Luisa Mesquita Pereira, Francois-Xavier Ricaut, Francesca Brisighelli, Irene Cardinali, Viola Grugni, Hovirag Lancioni, Vincenzo Lorenzo Pascali, Antonio Torroni, Ornella Semino, Giuseppe Matullo, Alessandro Achilli, Anna Olivieri, and Cristian Capelli. Population structure of modern-day italians reveals patterns of ancient and archaic ancestries in southern europe, 2018.

[7] Russell Thornton. *American Indian Holocaust and Survival: A Population History Since 1492.* University of Oklahoma Press, 1990.
